# Benchmarking GROMACS on Optimized Colab Processors and the Flexibility of Cloud Computing for Molecular Dynamics

**DOI:** 10.1101/2024.11.14.623563

**Authors:** Taner Karagöl, Alper Karagöl

## Abstract

<u>M</u>olecular <u>d</u>ynamics (MD) simulations are widely used computational tools in chemical and biological sciences. For these simulations, GROMACS is a popular open-source alternative among molecular dynamics simulation software designed for biochemical molecules. In addition to software, these simulations traditionally relied on costly infrastructure like supercomputers or clusters for <u>H</u>igh-<u>P</u>erformance <u>C</u>omputing (HPC). In recent years, there has been a significant shift towards using commercial cloud providers’ computing resources, in general. This shift is driven by the flexibility and accessibility these platforms offer, irrespective of an organization’s financial capacity. Many commercial compute platforms such as <u>G</u>oogle <u>C</u>ompute <u>E</u>ngine (GCE) and <u>A</u>mazon <u>W</u>eb <u>S</u>ervices (AWS) provide scalable computing infrastructure. An alternative to these platforms is Google Colab, a cloud-based platform, provides a convenient computing solution by offering GPU and TPU resources that can be utilized for scientific computing. The accessibility of Colab makes it easier for a wider audience to conduct computational tasks without needing specialized hardware or otherwise costly infrastructure. However, running GROMACS on Colab also comes with limitations. Google Colab imposes usage restrictions, such as time limits for continuous sessions, capped at several hours, and limits on the availability of high-performance GPUs. Users may also face disruptions due to session timeouts or hardware availability constraints, which can be challenging for large or long-running molecular simulations. We have significantly enhanced the performance of GROMACS on Google Colab by re-compiling the software, compared to its default pre-compiled version. We also present a method for integrating Google Drive to save and resume interrupted simulations, ensuring that users can secure files after session-timeouts. Additionally, we detail the setup and utilization of the CUDA and MPI environment in Colab to enhance GROMACS performance. Finally, we compare the efficiency of CUDA-enabled GPUs with Google’s TPUv2 units, highlighting the trade-offs of each platform for molecular dynamics simulations. This work equips researchers, students, and educators with practical MD tools while providing insights to optimize their simulations within the Colab environment.

## Introduction

<u>M</u>olecular <u>d</u>ynamics (MD) simulations have become a fundamental element in computational scientific/biological research, enabling scientists and researchers to explore and analyze the subjected atoms and molecules over a timed trajectory. These simulations are invaluable for testing various phenomena in biochemical sciences, such as drug interactions, protein folding, and the mechanical properties of nanomaterials [1, 2]. These MD simulations provide a dynamic and high-resolution view of molecular systems that is often unattainable through traditional experimental approaches [3]. However, the computational demands of these simulations are substantial, often requiring computational expertise and advanced algorithms to achieve meaningful results [4].

In recent years, the proliferation of cloud computing has improved how researchers access computational resources and their research outcome [5]. Cloud platforms, including <u>G</u>oogle <u>C</u>ompute <u>E</u>ngine (GCE) and <u>A</u>mazon <u>W</u>eb <u>S</u>ervices (AWS), offer flexible and scalable infrastructure that allows researchers to conduct experiments without the limitations imposed by traditional on-premises computing setups [6]. This shift resulted a more democratized access to <u>h</u>igh-<u>p</u>erformance <u>c</u>omputing (HPC), enabling a broader range of researchers utilize these resources for their scientific inquiries [7]. On the other hand, the computational costs and the required technical skills of these platforms may also impact accessibility and contribute to research inequalities, particularly in low-income settings.

Among these cloud solutions, Google Colab stands out as a particularly accessible platform for conducting scientific computations [8, 9, 10]. With its user-friendly interface, integration with Google Drive, and access to GPU and TPU resources, Colab allows users to engage in computational resources without the need for specialized hardware or significant financial investment. This accessibility is particularly beneficial for educational purposes, where students can experiment with MD simulations and gain hands-on experience in computational research.

Despite its many advantages, running GROMACS [11] (a widely used software package for MD simulations) on Google Colab comes with its own set of challenges. The platform, even with paid Pro+ subscriptions, imposes usage restrictions, such as session time limits that are typically capped at several hours and variable availability of high-performance GPUs. These constraints can significantly hinder the ability to perform large-scale simulations or to execute long-running calculations that require extended computational time. Furthermore, interruptions due to session timeouts can disrupt ongoing simulations, resulting in potential loss of data and requiring users to restart their computations from scratch. To address these challenges, this work focuses on enhancing the performance of GROMACS within the Google Colab environment. By re-compiling GROMACS and optimizing its configuration, we significantly improve its computational efficiency compared to default code, providing researchers with a more effective tool for their MD simulations. Additionally, we present a method for integrating Google Drive, enabling users to save their work and resume their simulations in a new session. This capability is crucial for ensuring that research can secure their files, as the loss of data may result in significant challenges in management or for the project.

Furthermore, we explore the setup and compilation of the CUDA and MPI environment within Colab to further enhance GROMACS performance. By integrating the power of GPUs, we aim to provide an understanding of how to optimize computational tasks effectively, which could be utilized by other scientific computing projects. In addition, we conduct a comparative analysis of CUDA-enabled GPUs versus Google’s TPUv2 units. This comparison not only informs users about the best computational strategies for their specific needs but also contributes to a broader understanding of utilizing computational resources in scientific research.

Ultimately, this study aims to equip researchers, students, and educators with the practical tools and insights necessary to optimize their simulations within the Google Colab environment. By addressing the limitations inherent in using GROMACS on cloud platforms, we hope to provide a greater accessibility and efficiency in molecular dynamics research, for educational use, innovative discoveries, and advancements in various scientific fields.

## Results

### GROMACS Availability Across Cloud Platforms

In this study, we assessed the availability of GROMACS on commercially accessible platforms, including <u>A</u>mazon <u>W</u>eb <u>S</u>ervices (AWS), Microsoft Azure, and <u>G</u>oogle <u>C</u>loud <u>E</u>ngine (GCE). All platforms were used with the latest Ubuntu distribution and were capable of installing GROMACS. AWS, Azure, and GCE offered SSH-based access [12], allowing remote users to manage simulations on these cloud environments. <u>M</u>olecular <u>d</u>ynamics (MD) simulations, depending on the system size, can take from several hours to days, thereby necessitating background execution. This allows simulations to continue even if the SSH session is interrupted. Data retrieval is another important process to conduct the analyses more efficiently, which was facilitated through SSH-based tools like WinSCP.

Most cloud services follow a pay-as-you-go model, highlighting the importance of setting usage limits to manage costs. In particular, we used 172-core Intel Sapphire Rapids CPUs to run simulations of up to 300 ns on the EAA1 protein [13], EAA2 and EAA3 [13], and large dimer structures [14]. Results indicate that cloud services, particularly GCE, provide a viable alternative to local computational resources for researchers, offering flexibility in selecting computing resources according to the scale of the project, thereby optimizing cost. Simplified instructions for using GROMACS on AWS and GCE are provided in the accompanying GitHub repository.

Google Colab Pro and Pro+ environments were also evaluated for their suitability for MD simulations. While Colab simplifies access by eliminating the need for SSH, the Pro version lacks support for background execution, a feature that is currently only available in the Pro+ environment. However, the Pro+ version has unclear time limits, generally up to 24 hours, necessitating frequent file backups to prevent data loss. Google Drive was used as an integrated file system to address this limitation, offering better data management within Colab. The availability of different processing options in Colab was further analyzed to determine the most suitable environment for MD simulations.

### Recompiling GROMACS for Colab’s Processing Units

In Colab, the default installation of GROMACS (version 2021) is without support for CUDA and MPI, both of which are crucial for optimizing performance [15, 16, 17, 18]. To enhance the simulation efficiency, we recompiled GROMACS (version 2024.3) with CUDA and MPI support on all available processor types. However, it is important to note that this recompilation process must be repeated for each new Colab session. Despite this inconvenience, performance benchmarks demonstrated significant improvements in simulation times after recompiling GROMACS.

### MPI Over-optimization for TPUv2

Another point to consider is that the TPU processing units are not yet fully optimized for GROMACS due to their recent development. In this study, we evaluated several MPI optimizations for TPUv2 tensor processing units. Without recompiling GROMACS in Colab, the initial test results for CHARMM-GUI 50ns performance with the melittin protein yielded a score of 29.598. Although this performance is double that of regular CPU usage, it is still relatively low given the TPU’s potential. When MPI was enabled, the score improved to 66.122, nearly doubling the initial result. However, GROMACS MPI support is not fully optimized for tensor processing units. To explore further, we tested several configurations involving oversubscription of the processing units (which is not recommended for general users). With 96 units, the software encountered multiple abort errors, and similar errors occurred with 90 oversubscribed processors. Ultimately, we tested with 72 processors, which successfully completed the tests without errors. The final performance score achieved was 110.267, nearly four times the initial score. These outcomes were also observed with the Max Planck Institute benchmark tools, particularly for membrane simulations, where performance increased from 12.433 with MPI to 53.406 with 72-core oversubscribed MPI.

### Benchmark Results and Performance Comparison

Performance tests were conducted using the benchmarking tools from the Max Planck Institute [5, 19, 20, 21] and custom simulations of a small melittin (from AlphaFold Database [22, 23], UniProt [24] ID: P01501) membrane protein created using CHARMM-GUI [25, 26, 27, 28, 29, 30, 31]. The results demonstrated that recompiling GROMACS with CUDA and MPI support substantially improved performance across different processor types. (Table 2)

**Table 1.** Selected benchmark results for custom melittin and Max Planck Institute benchmarking models across different Google Colab processor setups and compiler options.

| Processor Type |  | Benchmarking Type and Score |  |  |  |  |
| --- | --- | --- | --- | --- | --- | --- |
|  |  | Melittin<sup>1</sup> | Melittin-50ns<sup>2</sup> | MEM<sup>3</sup> | RIB<sup>4</sup> | PEP-h<sup>5</sup> |
| **A100** | A100-default | 12.736 | 13.363 | - | - | - |
|  | A100-CUDA | 322.718 | 342.727 | 45.58 | 7.258 | 1.954 |
| **L4** | L4-default | 12.759 | 13.384 | - | - | - |
|  | L4-CUDA | 322.597 | 342.640 | 33.391 | 6.484 | 0.822 |
| **T4** | T4-default | 9.307 | 9.756 | - | - | - |
|  | T4-CUDA | 123.526 | 125.991 | 45.751 | 3.625 | 0.437 |
| **TPUv2** | TPUv2-default | 28.033 | 29.598 | 12.433 | 2.374 | 0.244 |
|  | TPUv2-MPI | 62.551 | 66.122 | - | - | - |
|  | TPUv2-MPI-72 | 104.740 | 110.267 | 53.406 | 3.814 | 0.328 |
| **CPU** | CPU-default | 13.200 | 13.855 | 7.689 | 0.469 | 0.038 |

<sup>1,2</sup> Molecular dynamics simulations of the melittin protein (AlphaFold Database, UniProt ID: P01501) were conducted using input files from CHARMM-GUI. Full simulation results were calculated from two equilibration steps and two production steps (see Methods).
<sup>3,4,5</sup> Molecular dynamics benchmarks from the Max Planck Institute were used to assess simulation performance across various systems (see Methods).

We conducted tests across all available processor types on Google Colab, using various compilation options. In custom benchmarking of a small melittin membrane protein model generated with CHARMM-GUI, tests were conducted across two equilibration steps and two production steps (Figure 1), with the final score calculated based on the step counts. Initially, the default-mode TPUv2 provided the best performance without recompilation (28.033 performance score). However, after enabling CUDA support, GPUs outperformed all other processor types, achieving significantly faster simulation times. For example, A100 performance scores increased dramatically from 12.736 to 322.718—a nearly 25-fold improvement. Despite the A100 having more computational units, both A100 and L4 GPUs showed similar results (322.718 and 322.597 with CUDA respectively). Based on these findings, we recommend using CUDA-compiled versions on Colab’s A100, L4, and T4 GPUs to maximize simulation efficiency. Interestingly, TPUv2 processors, with MPI support and oversubscribed cores, showed considerable improvements (104.740) and performed comparably with T4 GPUs (Figure 2). For researchers prioritizing cost-effectiveness, T4 and L4 GPUs offer the best performance-to-cost ratio for CHARMM-GUI membrane simulations, although this may vary based on simulation type and pricing policy of Google Colab.

**Figure 1.**
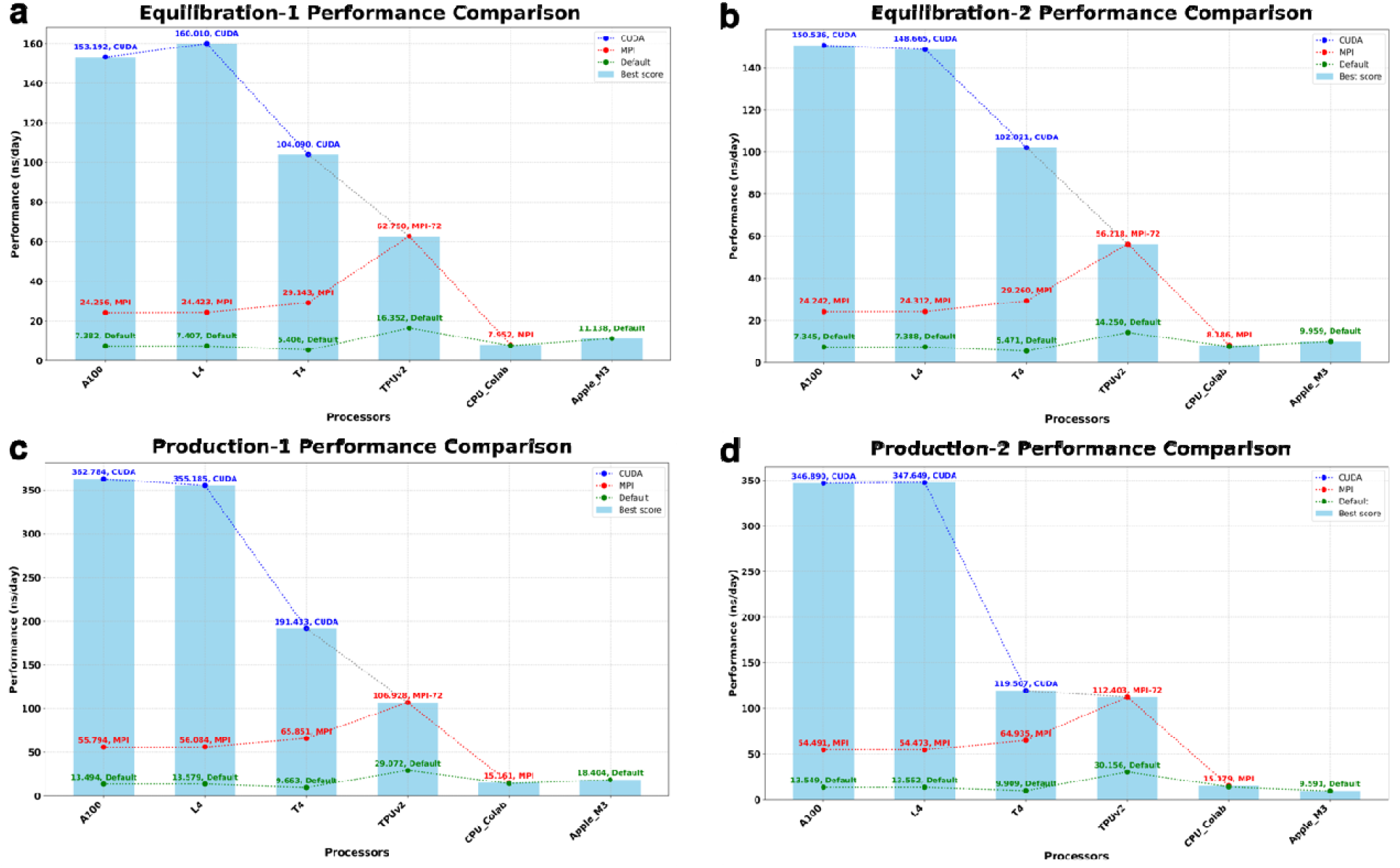
Custom benchmarking of a melittin membrane protein model evaluated across two equilibration steps and two production steps. Molecular dynamics simulations of the melittin protein (AlphaFold Database, UniProt ID: P01501) were conducted using input file from CHARMM-GUI. Performance metrics include comparisons across compiler options (blu line: CUDA, red line: MPI, green line: Default) and various processor setups (A100, L4, T4, TPUv2, CPU), alongside benchmarking results for the 2024 Apple M3 (8GB) model for **a, b)** two equilibration steps and **c, d)** two production steps. The best result for each processor is shown in light blue bars.

**Figure 2.**
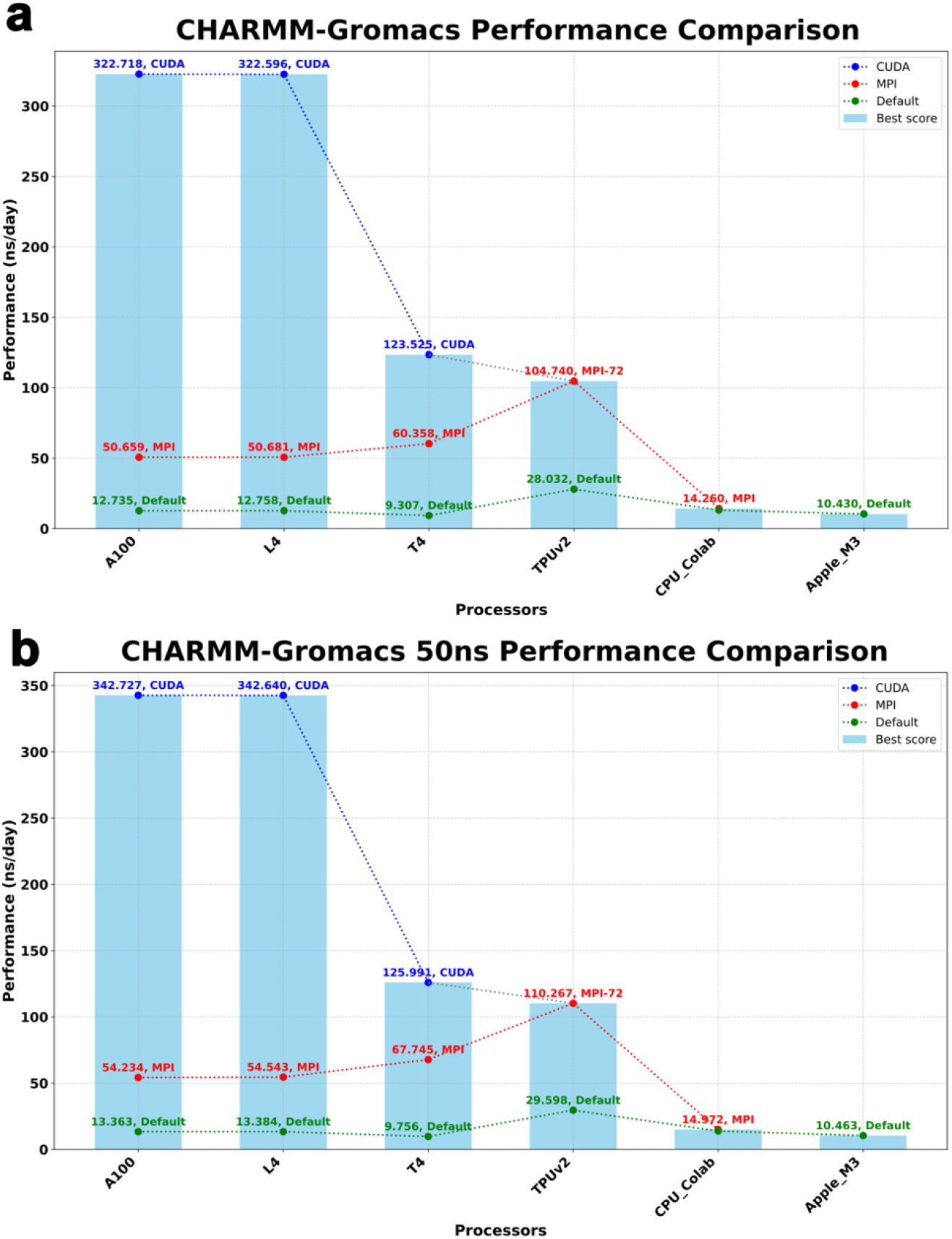
Custom benchmarking of a melittin membrane protein model evaluated for full simulation. Molecular dynamics simulations of the melittin protein (AlphaFold Database, UniProt ID: P01501) were conducted using input files from CHARMM-GUI. Performance metrics include comparisons across compiler options (blue line: CUDA, red line: MPI, green line: Default) and various processor setups (A100, L4, T4, TPUv2, CPU), along with benchmarking results for the 2024 Apple M3 (8GB) model. Full simulation results were calculated from two equilibration steps and two production steps, based on: **a)** the default Gromacs setup, and **b)** the 50ns configuration. The best result for each processor is shown in light blue bars.

To further evaluate our configurations, we used standard molecular dynamics benchmarks from the Max Planck Institute [5, 19, 20, 21] to assess benchmark times across different simulation types. Notably, the results revealed that the optimal processor configuration is highly dependent on the simulation type. For the membrane (MEM) benchmark, the over-optimized MPI on TPUv2 delivered the best performance, whereas the A100 GPU with CUDA outperformed other simulation types in general. However, for the ribosome (RIB) benchmark, the performance gap narrowed, suggesting that the A100 may not be the most cost-effective choice for this type of simulation. Protein (PEP-h) benchmarking results aligned closely with those from the CHARMM-GUI melittin protein membrane custom benchmarks, except higher performance of A100 GPUs (Figure 3).

**Figure 3.**
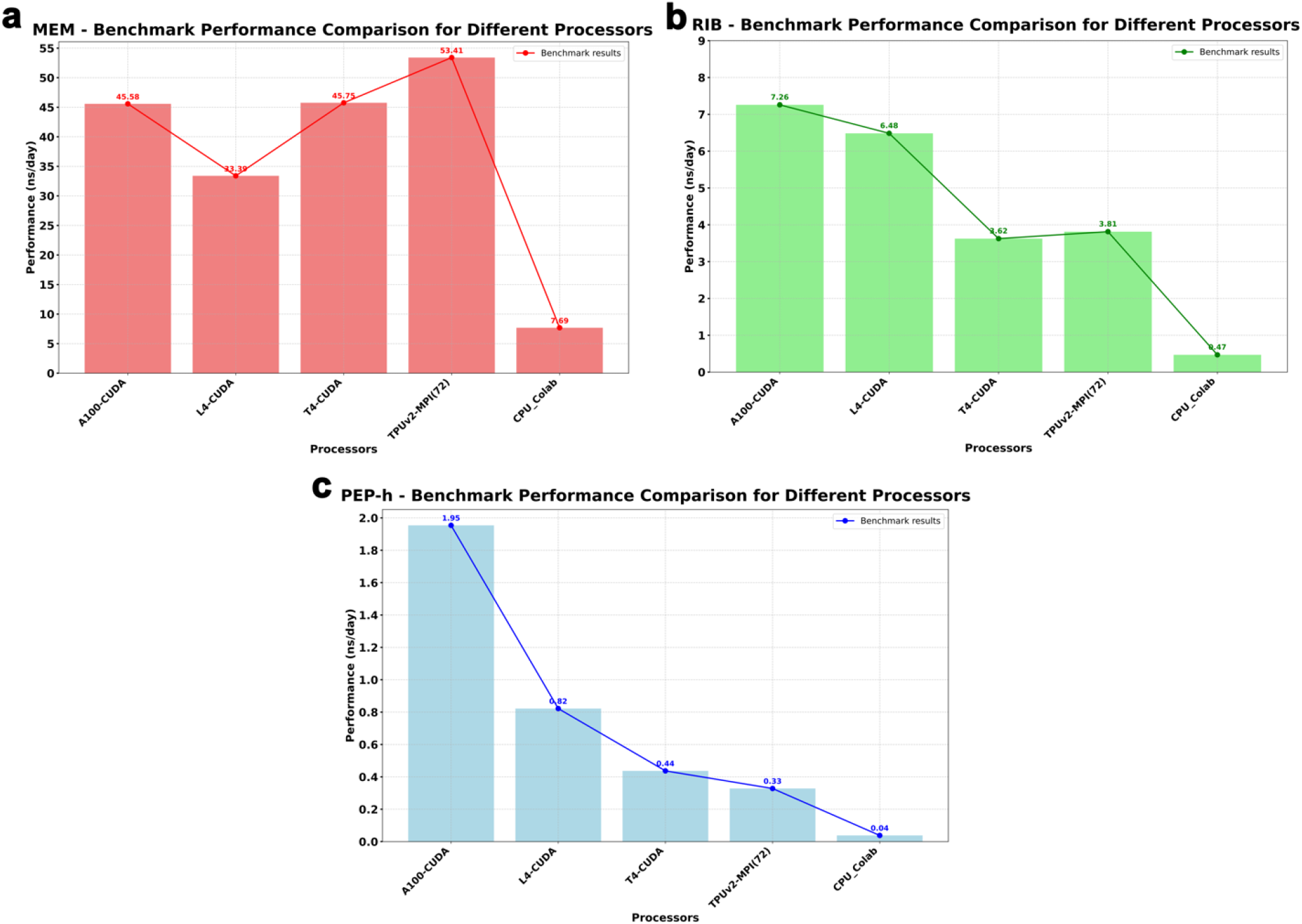
Standard benchmarking from the Max Planck Institute evaluated across different simulation types and processor configurations. Molecular dynamics benchmarks from the Max Planck Institute were used to assess simulation performance across various systems. Results show that the optimal processor configuration depends on the simulation type. **a)** For the membrane (MEM) benchmark TPUv2 with MPI performed best for, while A100 GPUs with CUDA excelled for other simulations. **b)** For the ribosome (RIB) benchmark, the performance gap narrowed. c) Protein (PEP-h) results showed that A100 GPUs provided higher performance.

Overall, GPUs equipped with Colab’s performance with CUDA-enabled GPUs may even surpass that of local CPU-based university systems, offering greater flexibility for MD research at a lower cost.

## Conclusion

Recent advancements in computational methods for biology, such as structural predictions and docking simulations, have made <u>m</u>olecular <u>d</u>ynamics (MD) simulations increasingly popular. Structural prediction tools, often accessible through cloud-based web interfaces, allow researchers worldwide to perform analyses regardless of local computational resources. However, limited access to <u>h</u>igh-<u>p</u>erformance <u>c</u>omputing (HPC) clusters remains a significant barrier for researchers in many low-income countries, where universities often struggle to pay the high costs of maintaining large-scale cluster systems. This study demonstrates that commercially available cloud solutions, such as Google Colab, can serve as cost-effective alternatives to local HPC resources. The recompilation process to achieve optimized GROMACS runs must be repeated for each new Colab session. Despite this inconvenience, performance benchmarks demonstrated significant improvements in simulation times after recompiling GROMACS. With recompiling, longer simulations in Colab become possible and more accessible. During simulations, we observed that there is a room for optimization of the GROMACS, especially for TPUs. Our benchmarking results further highlight the importance of software optimization for achieving efficient MD simulations in Google Colab, and its potential as a viable option for resource-constrained environments.

## Methods

### Accessibility and User-end Interfaces

We evaluated the accessibility of GROMACS across multiple cloud computing platforms, including <u>A</u>mazon <u>W</u>eb <u>S</u>ervices (AWS), Microsoft Azure, <u>G</u>oogle <u>C</u>loud <u>E</u>ngine (GCE), and Google Colab, all configured with the latest Ubuntu distribution. GCE, Azure, and AWS provided SSH-based remote access, allowing users to manage <u>m</u>olecular <u>d</u>ynamics (MD) simulations securely. In contrast, Google Colab offered a simplified, browser-based interface that eliminated the need for SSH, though limitations such as lack of background execution in the Pro version posed challenges. For Colab, Pro+ was enabled and Google Drive was integrated as a file system for secure, continuous data backup. WinSCP and similar SSH tools facilitated data transfer from cloud platforms post-simulation, and a GitHub repository was created with setup instructions for users, specifically detailing GROMACS setup on AWS and GCE.

### Re-compiling of GROMACS for Available Processors

The default installation of GROMACS on Google Colab lacks support for CUDA and MPI, both of which are essential for achieving optimal performance on GPU-accelerated processors. To leverage Colab’s hardware fully, we recompiled GROMACS with CUDA and MPI support in each new session, focusing on compatibility with available GPUs and TPUs. The recompilation process allowed us to activate essential optimizations for MD simulations. Colab’s GPU options, such as A100, L4, and T4, along with TPUv2 processors, were tested post-recompilation to determine their suitability for MD tasks, with a particular focus on performance improvements achieved through CUDA and MPI.

### Benchmarking Models

For benchmarking, we conducted MD simulations of the melittin (from AlphaFold Database [22, 23], UniProt [24] ID: P01501) protein, chosen for its relevance and computational load suitability for cloud-based testing. Input files were generated through CHARMM-GUI [25, 26, 27, 28, 29, 30, 31], which provided the necessary starting structures and parameters for GROMACS simulations. The benchmarking process included both standard MD benchmarks from the Max Planck Institute [5, 19, 20, 21] and custom simulations of the melittin protein, allowing us to compare computational efficiency across processors. GCE’s 172-core Intel Sapphire Rapids CPUs were also tested previously [13, 14]. The benchmarking we used from the Max Planck Institute included the following simulations: benchMEM, which involves 82k atoms with a protein in a membrane surrounded by water and a 2 fs time step; benchPEP-h, with 12 million atoms, peptides in water, a 2 fs time step, and hydrogen bonds constrained; and benchRIB, which features 2 million atoms with a ribosome in water and a 4 fs time step.

### Benchmark Analysis and Visualization

Python scripts were used to analyze simulation performance data and generate benchmark graphs, providing visual insights into the performance differences across platforms. Python’s data processing libraries facilitated statistical analysis of simulation times across Colab’s processor types, enabling direct comparison between GPU and TPU performance with and without CUDA and MPI support. Benchmarking graphs, visualized through Python’s Matplotlib [32] libraries, illustrated the computational efficiency of Colab’s A100, L4, and T4 GPUs, as well as TPUv2 units, highlighting the performance boost observed post-recompilation with CUDA and MPI.

### Availability

The resource codes and usage instructions are freely available for academic and non-commercial use. They are provided as-is, without any advice, recommendations, or guarantees, and can be accessed at: https://github.com/karagol-taner/Molecular-Dynamics-on-AWS-and-Cloud-Computing

## Supplementary Information

The benchmark scores are available in the Supplementary Information as the file Supplementary_Information.xlsx.

## Acknowledgments

We thank the research community for valuable feedback during the testing.

